# Astrocyte immunosuppressive activity in glioblastoma depends on ZEB1 and is counteracted by CXCL14

**DOI:** 10.64898/2026.05.10.724074

**Authors:** Mathew Clement, Alex Gibbs, Ayesha Begum, Dorit Siebzehnrubl, Suresh Kaushik, Niharika Singh, Bhavana Gupta, Vasileios Eftychidis, Florian A. Siebzehnrubl

## Abstract

Glioblastomas are incurable and lethal brain cancers. Immunotherapies offer new and promising treatment options for glioblastoma patients, but the highly immunosuppressive nature of these cancers presents a challenging clinical obstacle. Glioblastoma immune evasion is driven by cell-cell interactions in the tumor microenvironment and recent studies have identified astrocytes as important contributors to immune silencing [1, 2]. Cell plasticity is a key feature of reactive astrocytes that drives heterogeneous, pro- or anti-inflammatory states [3], but the molecular regulators of astrocyte-immune interactions remain incompletely understood. Here, we investigate whether cell plasticity of glioblastoma-associated astrocytes promotes or opposes tumor progression and show that loss of astrocyte plasticity results in T-cell recruitment and immune activation. We evaluate how astrocytic cell plasticity contributes to immune functions in the glioblastoma microenvironment using single cell sequencing from preclinical models, in vivo genetic perturbations and in vitro mouse and human experimental systems. We show that astrocytes surrounding glioblastoma express the stem cell-associated transcription factor, ZEB1, and that conditional-inducible astrocytic deletion of *Zeb1* remarkably reduces glioblastoma growth and extends survival. Increased recruitment and activation of T cells in astrocytic *Zeb1*-deficient mouse models is linked to increased expression of the immunoattractant cytokine CXCL14, and viral delivery of CXCL14 in experimental glioblastoma models increases survival. Our data support that CXCL14 is a candidate therapeutic target for reprogramming the tumor microenvironment that can restrict and reduce glioblastoma growth and progression.

## Introduction

Glioblastoma (GBM) is an incurable disease with median survival rates of approximately 18-24 months with therapy [4]. Poor outcome is caused by diffusely infiltrative GBM cells that remain after resection and resist chemo-/radiotherapy [5]. Extensive tumor heterogeneity between patients [6], on the single cell [7, 8] and on the spatial level [9] contributes to GBM therapy resistance. Cell-cell interactions between heterogeneous GBM cells and the diverse cell types in the tumor microenvironment (TME) further add to the complexity of this disease [10]. For example, GBM immune evasion is driven by interactions with immune cells and microglia [11]. Manipulation of these interactions and improving immune cell function, particularly T cell function, therefore represents an attractive treatment strategy. Many studies and clinical trials are actively exploring ways to unlock the full killing capacity of CD8^+^ T cells that includes the use of immune checkpoint inhibitors, chimeric antigen receptor T cells (CAR T cells), bispecific antibodies, oncolytic viruses, tumor-infiltrating lymphocytes (TILs), combination therapies and vaccines (reviewed in [12] and [13]). The success of such treatments is reliant upon the presence of cytotoxic CD8^+^ T cells within the TME with increased prevalence in high-grade gliomas as compared to low-grade gliomas [11] However, these typically “cold” tumors are coupled with an immunosuppressive TME that promotes the tumor to grow unchecked. Therefore, a deeper understanding of the mechanisms that control T cell functionality within the TME may be exploited to effectively stimulate a robust anti-cancer T cell response within the brain.

Gene expression signatures indicate that GBM-associated astrocytes (GAA) contribute to anti-inflammatory signaling in the TME and recent studies have shown that GAA are key modulators of immune function and important contributors to GBM immune evasion [1, 2, 14, 15]. Yet, the molecular regulators governing GAA functions and immune regulatory phenotypes remain mostly obscure. Gene expression profiling has shown that GAA are characterized by increased cell plasticity [1, 16]. For example, GAA exhibit increased expression of stem cell-associated cytoskeletal proteins VIM and NES, even compared to reactive astrocytes in other diseases [1, 17]. Therefore, it is likely that the GBM TME reprograms astrocytes to a distinct cell state and that this reprogramming depends on transcriptional regulators of cell plasticity.

Here, we identify the transcription factor ZEB1 as a regulator of cell plasticity in GAA that controls astrocyte-immune interactions and is necessary for astrocytes to generate a pro-tumorigenic environment. Genetic deletion of *Zeb1* reverts host astrocytes to an anti-tumorigenic and pro-inflammatory phenotype that results in increased activation of T cells which promotes immune clearance of experimental GBM. Mechanistically, we identify the chemokine CXCL14 as downstream target of ZEB1. Increased CXCL14 expressing after *Zeb1* loss or through viral delivery causes increased T cell infiltration into the brain and T cell activation. Our work identifies a previously unrecognized role for ZEB1 in the brain and provides proof-of-concept that CXCL14 may be therapeutically beneficial in GBM.

## Results

We have previously shown that ZEB1 regulates cell plasticity in GBM [18] and in neural stem cells, where *Zeb1* loss also reduces astroglial differentiation [19]. Many studies have documented that ZEB1 is not only present in cancer cells but also within the TME, for example in cancer-associated fibroblasts [20]. To investigate how ZEB1 can modulate astrocyte and immune cell function in the GBM TME, we probed publicly available single cell RNA-seq (scRNAseq) datasets and performed immunofluorescence staining.

### Zeb1 is expressed in GAA surrounding GBM

Only few publicly available GBM scRNAseq datasets contain significant numbers of host cell populations. We used extended GBmap [21] to evaluate ZEB1 expression across different cell types. When comparing normalized read counts, we found a comparable range of ZEB1 expression levels in oligodendrocyte precursor cells (OPCs), GBM cells, neurons, and astrocytes, as well as in vascular endothelia and mural cells, and microglia, mature T cells, and macrophages, with other cell types showing considerably lower ZEB1 expression (Fig. 1A, S1A,B). To validate ZEB1 expression in neural cell types, we used immunofluorescence staining in the mouse brain and detected ZEB1 in all astrocytes (Fig. 1B), but not in neurons (Fig. 1C), or oligodendrocytes (Fig. 1D). Since GBmap is a human dataset, the absence of ZEB1 protein in murine neurons may be due to species differences or other, post-translational mechanisms. OPCs showed a heterogeneous staining pattern with only some OPCs co-expressing ZEB1 (Fig. S1C). We observed ZEB1 expression in some endothelial cells, but apparently at lower levels than in astrocytes (Fig. S1D). Next, we investigated the TME of GBM orthotopic xenografts and found that ZEB1 is frequently expressed in GAA (Fig. 1E). Quantification of ZEB1 in GAA across two different xenograft models revealed that the percentage of ZEB1-positive astrocytes is heterogeneous (Fig. 1F). Immunofluorescence staining of human patient GBM tissue validated ZEB1 expression in GAA (Fig. 1G). These findings prompted us to ask what the key functions of ZEB1 in GAA are.

**Figure 1:**
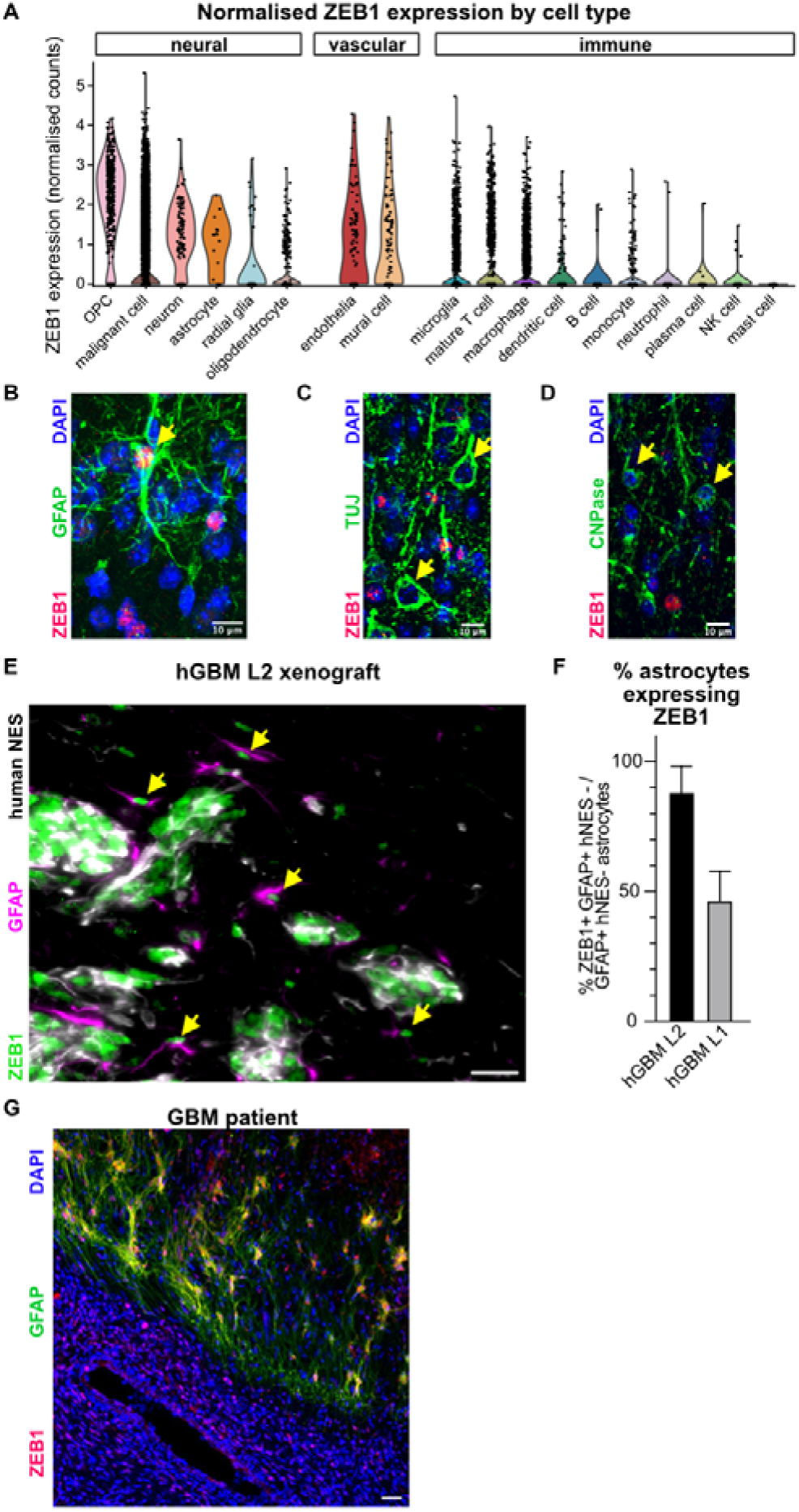
Zeb1 is expressed in GAA. **(A)** Quantification of ZEB1 expression in scRNAseq data from human GBM patients (Extended GBmap, [21]). In the mouse brain, ZEB1 protein expression is detectable in astrocytes (co-stained with GFAP, **B**), but not in neurons (co-stained with bIII-tubulin, TuJ, **C**) or oligodendrocytes (co-stained with CNPase, **D**). Images from three independent mice showed similar results. See Figure S1 for additional images. **(E)** Representative image from a patient-derived GBM xenograft showing ZEB1 expression in GAA (GFAP+/human-specific Nestin-, arrows). Images from three independent mice showed similar results. **(F)** Quantification of ZEB1 expression GAA in xenografted patient-derived GBM lines (hGBM L2 and hGBM L1); n = 5 areas per brain from 2 independent transplants (hGBM L2) and 3 areas from 3 independent transplants (hGBM L1). Data represent mean ± s.e.m. **(G)** Representative image from a GBM patient showing ZEB1 expression in GAA (GFAP+) at the tumor-brain interface. Scale bars: B-D 10 µm; E, G 20 µm.

### Genetic deletion of Zeb1 in host astrocytes reduces tumor growth and invasion

To investigate ZEB1 functions in the GBM TME, we used a transgenic mouse model for conditional-inducible deletion of *Zeb1* in astrocytes (Fig. 2A, [19]). GLAST::CreERT^2^,Zeb1^f/f^,R26-loxP-Stop-loxP-tdTomato mice (Zeb1 KO hereafter) allow targeted ablation of *Zeb1* in astrocytes and lineage tracing upon Tamoxifen administration, with GLAST::CreERT^2^,R26-loxP-Stop-loxP-tdTomato mice serving as lineage-traceable controls (controls hereafter). To investigate whether *Zeb1* loss in host astrocytes affected GBM histopathology, we induced genetic recombination by Tamoxifen administration and intracranially implanted GFP-labelled murine KR158B GBM cells one week later. Four weeks after tumor implantation we examined tumor size and invasion (Fig. 2B). Immunofluorescence staining confirmed successful ablation of ZEB1 in GAA (Fig. 2C). We then evaluated brains from control and Zeb1 KO mice and found that tumors in the Zeb1 KO group showed smaller volumes and less invasion into the surrounding tissue than in control animals (Fig. 2D-F). Noting that GAA in the Zeb1 KO group were morphologically different from control astrocytes, we performed immunofluorescence staining for the astrocytic marker GFAP. This revealed that control astrocytes in the GBM TME had more hypertrophic morphology and increased GFAP levels compared to Zeb1 KO astrocytes (Fig. 2G). This is consistent with decreased astrocyte reactivity in tumor-bearing Zeb1 KO mice. Recent work has shown that GBM-instructed astrocytes can suppress immune cell activation in the GBM TME [2, 15]. Because reactive astrocytes are known to interact with, and affect the activation of, immune cells in the brain, we next asked whether *Zeb1* deletion results in altered immune activation.

**Figure 2:**
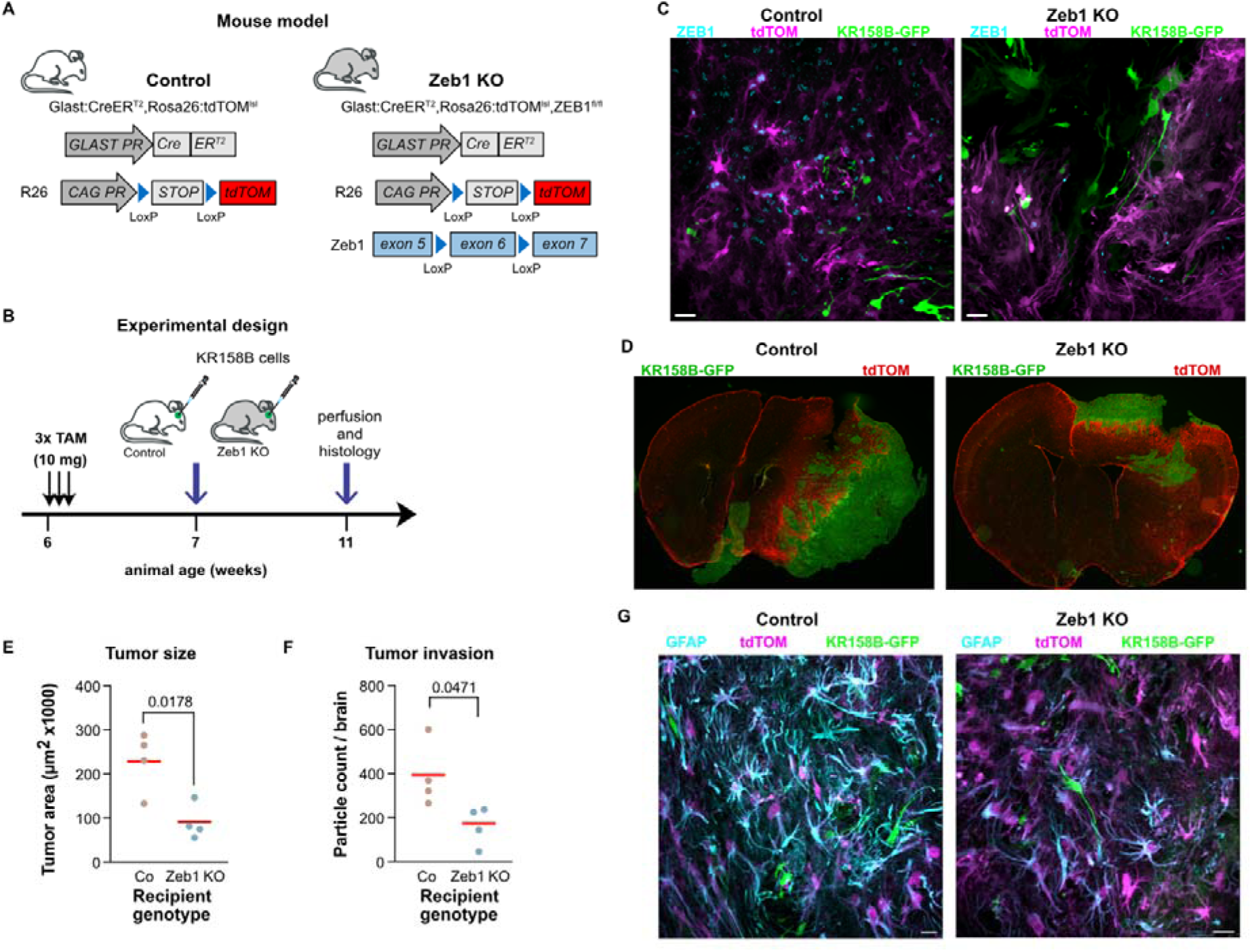
Genetic deletion of Zeb1 in host astrocytes reduces tumor growth and invasion. **(A)** Transgenic mouse model for conditional-inducible deletion of Zeb1 in astrocytes including a tdTomato reporter. **(B)** Experimental design. Recombination was induced 1 week prior to intracranial implantation of invasive murine KR158B GBM cells and brains were assessed 4 weeks later. **(C)** Representative images (single confocal planes) of GAA, demonstrating successful deletion of Zeb1 in ZEB1 KO but not control animals. Images from three independent mice showed similar results. **(D)** Representative images showing reduced tumor size and invasion in ZEB1 KO mice compared to controls. Images from four independent mice showed similar results. **(E,F)** Quantification of tumor area (E) and invasion (F) across multiple brain sections (averages for n=4 mice per genotype, Welsh’s t test, red line shows mean). **(G)** Representative images (single confocal planes) showing loss of GFAP expression in ZEB1 KO astrocytes compared to controls. Images from three independent mice showed similar results. Scale bars: C, G 20 µm.

### Increased survival following genetic targeting of Zeb1 in an immune evasive GBM model

To test this, we utilized a syngeneic murine GBM model that is capable of immune evasion (NPE-IE cells, [22]). We implanted NPE-IE cells orthotopically into recipient control or Zeb1 KO mice and evaluated overall survival, as well as single cell transcription profiles (Fig. 3A). Orthotopic implantation of NPE-IE cells into control mice resulted in malignant brain tumors in 5/8 mice with a median survival of 38 days, similar to what was reported before for these cells [22]. By contrast, NPE-IE grafts into Zeb1 KO mice formed tumors in only 1/11 mice with the remaining animals surviving >70 days (Fig. 3B; a control cohort of wild type mice implanted in the same experiment showed 2/4 mice forming tumors with a median survival of 47.5 days). Histological analysis showed a smaller tumor in Zeb1 KO mice which did not cause noticeable midline shift (Fig. 3C). We observed traces of GFP+ punctae in microglia in tissue of surviving Zeb1 KO mice grafted with NPE-IE cells (data not shown) and therefore speculated that immune cell activation led to clearance of grafted tumor cells in Zeb1 KO mice. To validate this hypothesis, we performed single cell RNAseq from NPE-IE grafted control and Zeb1 KO mice at an earlier timepoint using the 10X Genomics platform (Fig. 3A, S2A). We identified 12 distinguishable cell clusters using the Seurat analysis pipeline and used Enrichr on cluster-specific markers to help identify cell types, which revealed 2 separate clusters of astrocytes (termed astrocyte_1 and astrocyte_2), as well as immune cells (macrophages, microglia, and T cells) and NPE-IE cells (termed GBM; Fig. 3D). Analysis of top 50 differentially expressed genes within each cluster confirmed a clear separation of cluster identities (Fig. 3E). We found copy number variations (using inferCNV) only in the GBM cluster (Fig. 3F), but not in any of the other cell clusters (Fig. S2B). Of note, copy number variations in GBM cells included common genomic alterations associated with GBM, such as *Pten* deletion and amplification of *Pik3r1*, *Pik3ca*, *Akt1*, *Sox2*, and *Irs2*. Analysis of cluster-specific KEGG pathways revealed enrichment of immune signaling pathways in immune cell clusters (e.g., T cell receptor signaling, Antigen processing and presentation, Cytokine/receptor interaction), proliferative signatures in GBM cells (e.g., DNA replication, cell cycle), and metabolic signatures in astrocytes (e.g., OxPhos, Biosynthesis of unsaturated fatty acids, PPAR signaling; Fig. 3G). These results support correct cell type allocation to Seurat clusters.

**Figure 3:**
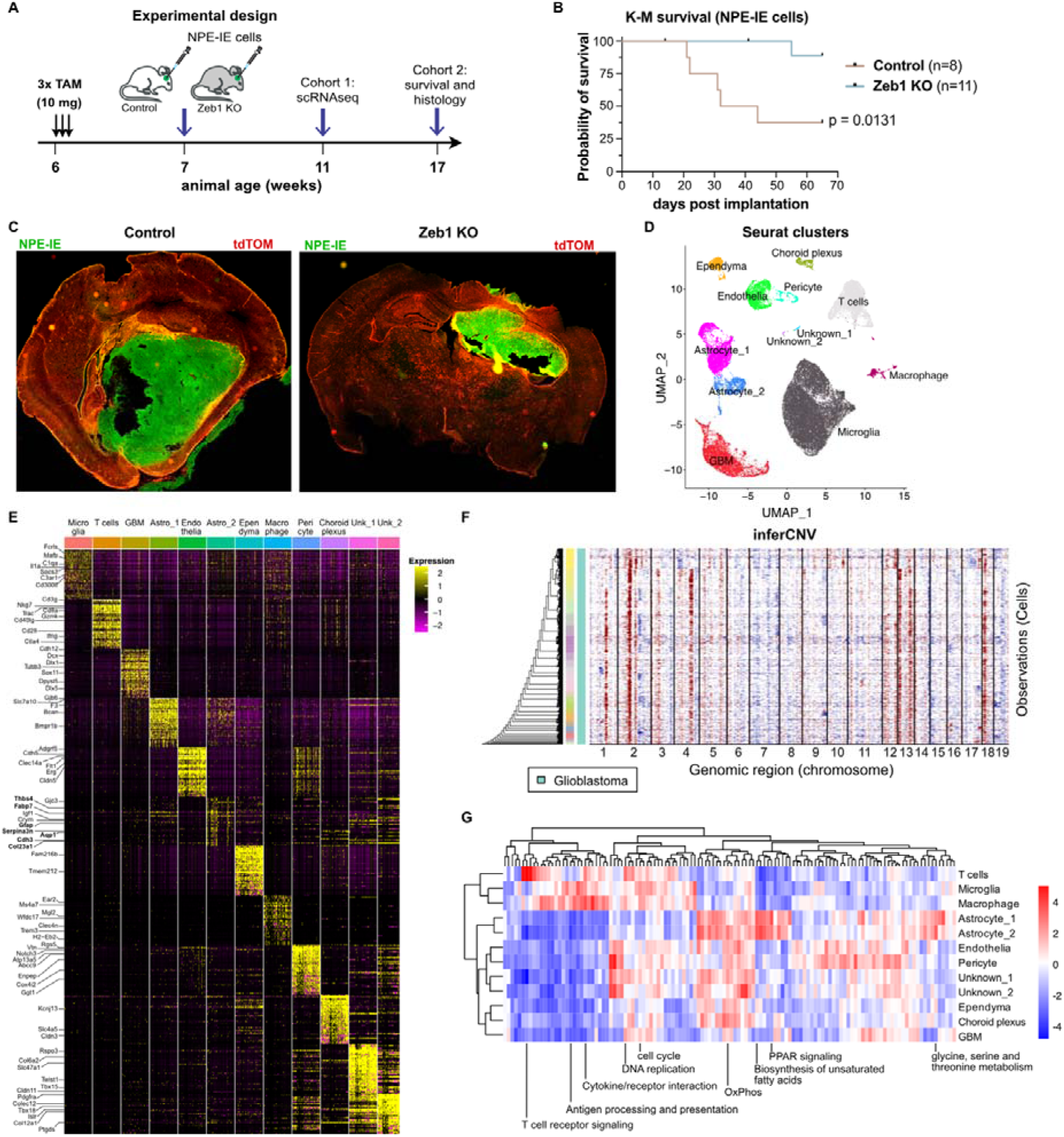
Genetic targeting of Zeb1 in an immune evasive GBM model increases survival. **(A)** Experimental design. Recombination was induced 1 week prior to intracranial implantation of immune evasive murine NPE-IE GBM cells. Tissue was harvested from some animals after 4 weeks for scRNAseq profiling (Cohort 1), while others were culled at end point to evaluate histology (Cohort 2). **(B)** Kaplan-Meier survival curve shows significant increase in survival of Zeb1 KO mice. Note that NPE-IE cells generate tumors in 50-60% of recipients. Log-Rank test. **(C)** Representative images of NPE-IE tumors in control and Zeb1 KO mice. Similar results were observed in 5 mice (control) and 2 mice (Zeb1 KO). **(D)** Annotated UMAP of Seurat clusters identifying different cell types in combined scRNAseq data from control and Zeb1 KO mice. **(E)** Heatmap showing top 50 DEG for each cluster. Key cell-type associated genes are indicated. **(F)** CNV plot from inferCNV showing substantial genomic aberrations (duplications in red, deletions in blue) in the GBM cell cluster. See Figure S2 for CNV plot of other clusters. **(G)** Heatmap showing differentially enriched KEGG pathways associated with specific clusters.

### scRNAseq reveals gene expression changes related to immune activation in Zeb1 KO astrocytes

To evaluate cell-type specific differences of astrocytic Zeb1 deletion, we compared single cell RNAseq data from control and Zeb1 KO mice. Overall clustering was comparable between control and Zeb1 KO cells, with only minor changes in cell numbers across clusters (Fig. 4A). When comparing *Zeb1* transcripts between control and Zeb1 KO mice, we found a strong reduction in the astrocyte_2 and a weaker but significant loss in the astrocyte_1 cluster, while GBM cells showed no change in *Zeb1* levels (Fig. 4B, S2C). Concomitantly, the astrocyte_2 cluster shows strongest differential gene expression between control and Zeb1 KO animals (Fig. 4C), yet we identified differential gene expression also in the astrocyte_1 cluster (Fig. 4D) and in GBM cells (Fig. 4E). Cells in the astrocyte_2 cluster showed heterogeneity in the expression of astrocyte subtype signatures, with *Zeb1* deletion promoting signatures of mature (AST2, [23]) and reactive (A2, [24]) subtypes (Fig. S3). Enrichment analysis of gene ontologies in astrocyte_2 cells identified reduction in ontologies associated with neurodevelopment and migration and increased metabolic signatures, in line with expected functions of *Zeb1* (Fig. 4F, S4A,B) [19, 25–27]. Notably, we also observed a significant reduction in immune signatures, which prompted us to investigate potential interactions between astrocytes and immune cells, as astrocyte-immune interactions have recently garnered attention as contributors to an immunosuppressive environment in GBM [2, 15]. We used LIANA [28] to evaluate ligand-receptor interactions, specifically focusing on astrocyte_2 as a source (Fig. S4C). While astrocytes interacted with immune cells in control animals (Fig. 4G), the LIANA cross-database consensus identified an exclusive interaction between astrocyte_2 and T cells in Zeb1 KO mice, which included astrocytic *Cxcl14* and *Cxcr4* on T cells (Fig. 4H, S4D,E). Pseudotime analysis further validated that gene expression changes in Zeb1 KO animals in the astrocyte_2 cluster preceded changes in immune cell clusters, with GBM cells affected last (Fig. 4I). These data support that Zeb1 loss results in altered astrocyte gene expression and signaling to T cells mediated via a Cxcl14 to Cxcr4 interaction.

**Figure 4:**
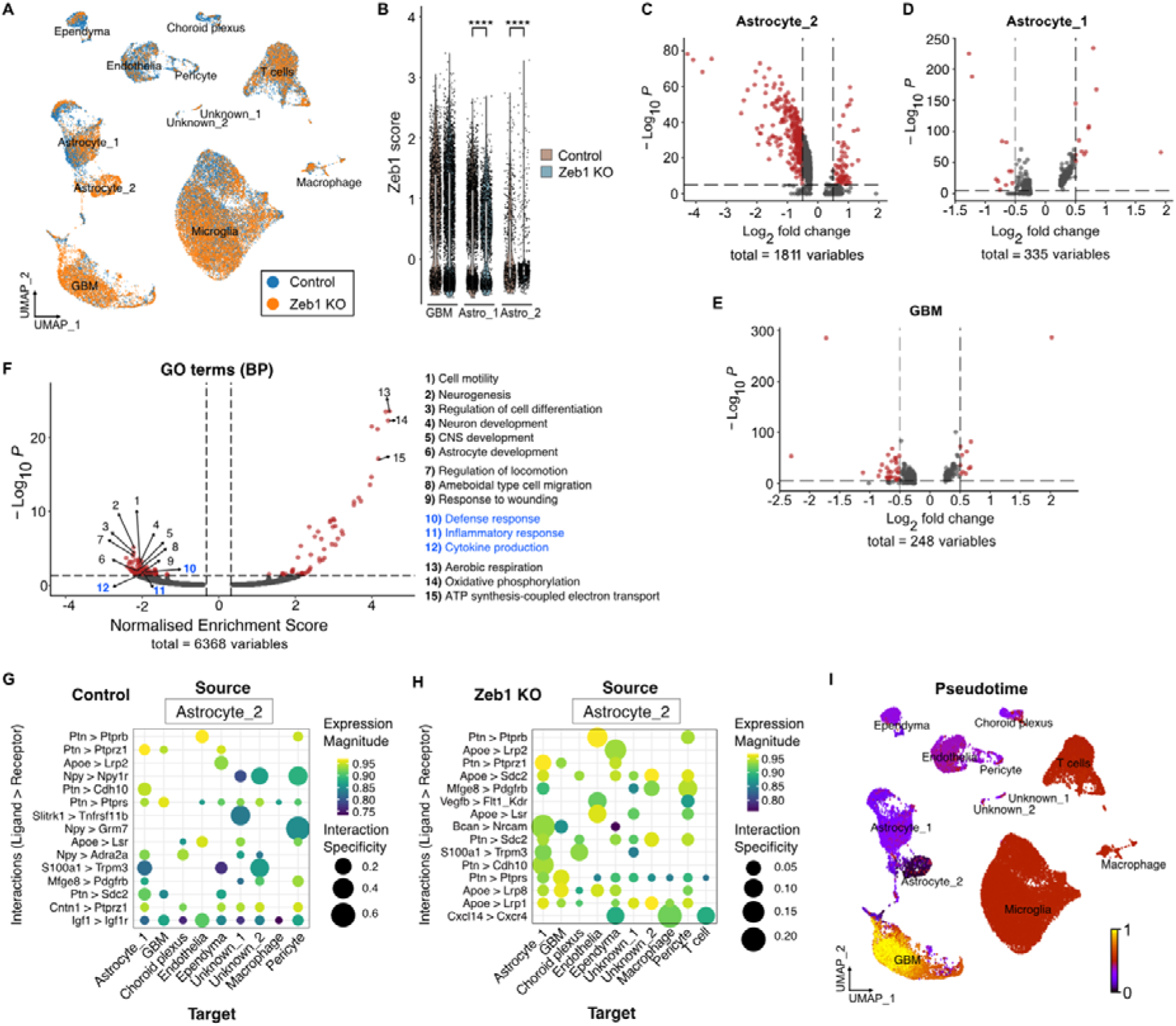
scRNAseq reveals gene expression changes related to immune activation in Zeb1 KO astrocytes. **(A)** UMAP plot shows similar proportions of clusters in control and Zeb1 KO mice. **(B)** Violin plot showing reduced Zeb1 expression in astrocyte_2 and astrocyte_1 clusters, but not in GBM cells. **(C-E)** Volcano plots showing DEGs in astrocyte_2 (C), astrocyte_1 (D), and GBM (E) clusters (P < 0.05, log_2_FC > 0.5). **(F)** GSEA enrichment plot for gene ontologies (GO BP) in astrocyte_2 cluster (Zeb1 KO vs control). Key GO terms are highlighted. Note changes in immune-related signatures (10-12; P < 0.05, normalized enrichment score > 0.5). LIANA ligand interaction dot plots for immune-related signatures changed in control **(G)** vs Zeb1 KO **(H)** astrocyte_2 cluster. **(I)** RNA velocity pseudotime plot showing initial gene expression changes in astrocyte_2 cluster, followed by later changes in immune cell clusters and finally GBM cells.

### Zeb1 KO in GAA causes site-specific immune cell activation

We next analyzed immune cell-specific gene expression changes in Zeb1 KO mice, finding only minor changes in differential gene expression of T cells (Fig. 5A) and microglia (Fig. 5B). Nevertheless, over-representation analysis of differentially expressed genes in T cells revealed significant enrichment for gene ontologies (BP) associated with immune activation (Fig. 5C). Comparative analysis of gene expression signatures of T cell subtypes showed a general increase in all signatures in Zeb1 KO mice (Fig. 5D). To validate potential changes in T cell activation in Zeb1 KO mice, we performed flow cytometry from processed brains and peripheral blood of NPE-IE implanted control and Zeb1 KO mice. This revealed an increase in overall T cell number, as well as significantly elevated IFNy+ and CD69+ activated cytotoxic T cells in the brain of Zeb1 KO animals (Fig. 5E, S5). By contrast, there was no change in total T cell numbers in the peripheral blood, but a small increase in CD69+ activated T cells (Fig. 5F). Similarly, overall CD4+ T cell numbers and IFNy+ and CD69+ activated CD4+ T cells were significantly elevated in the brain of Zeb1 KO animals (Fig. 5G) but remained unchanged in peripheral blood (Fig. 5H). This strongly supports site-specific, increased activation of cytotoxic T cells following astrocytic Zeb1 deletion, which may explain the remarkable increase in survival of tumor-implanted Zeb1 KO mice.

**Figure 5:**
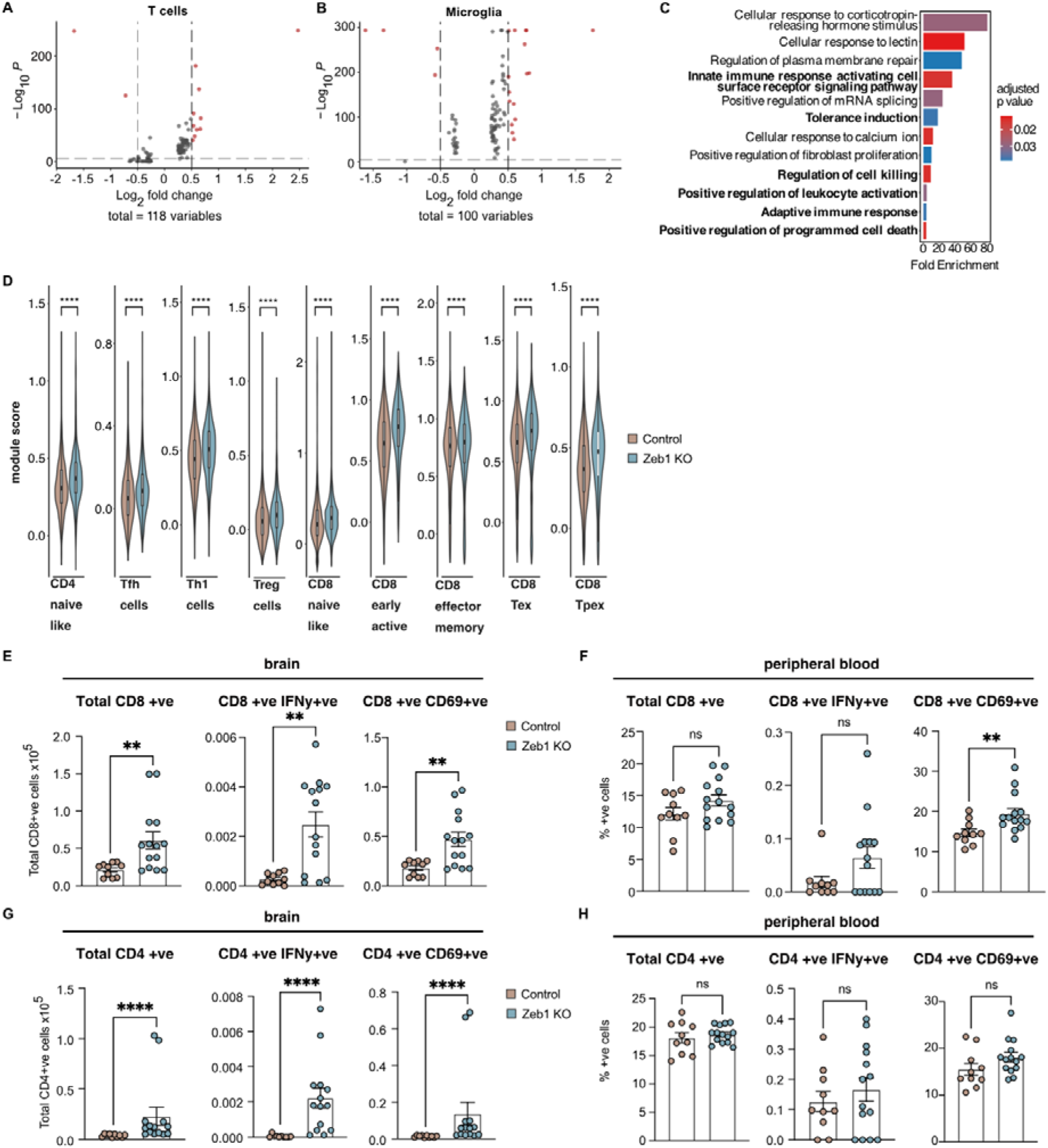
Zeb1 KO in GAA causes site-specific immune cell activation. **(A, B)** Volcano plots showing DEGs in T cell (A) and microglia (B) clusters (P < 0.05, log_2_FC > 0.5). **(C)** Bar plot showing ORA for GO terms (BP) in T cells. Key terms related to immune activation are highlighted. **(D)** Violin plots showing enrichment of module scores for T cell subtypes in Zeb1 KO vs control mice. **(E-H)** Flow cytometric analysis of CD8+ (E,F) and CD4+ (G,H) T cells in the brain (E,G) and peripheral blood (F,H) of Zeb1 KO and control mice (n = 5 mice for control and 7 mice for Zeb1 KO; 2 replicate experiments per animal plotted per graph). Zeb1 KO mice show increased numbers or total T cells as well as increased activation of CD8+ and CD4+ T cells indicated by co-expression of IFNy or CD69. Contrastingly, T cells from peripheral blood are unchanged. Mann-Whitney test; data represent mean ± s.e.m.

### scRNAseq analysis of human GBM patient datasets reveals signatures of immune activation in GAA stratified for ZEB1 expression

To validate our findings in human datasets, we curated a set of GAA from publicly available single cell RNAseq datasets at CELLxGENE [29] (Fig. 6A). We stratified GAA into quartiles based on ZEB1 expression (Fig. 6B). Differential gene expression analysis revealed a large number of differentially expressed transcripts between ZEB1-highest and ZEB1-lowest astrocyte quartiles (Fig. 6C), which showed enrichment of several gene ontologies associated with immune function (Fig. 6D). We then compared differentially expressed transcripts between ZEB1-deficient human and mouse datasets and found 340 overlapping genes across both sets (Fig. 6E). Subsetting this conserved gene set into candidates that were dysregulated in the same direction (Fig. 6F) revealed 19 candidates which were upregulated in both human and murine ZEB1-deficient astrocytes, of which 5 overlapped with immune-related gene ontologies (*GAPDH*, *HSPA8*, *CLU*, *CXCL14*, *APOE*; Fig. 6G). Because we identified CXCL14 as candidate ligand in cell-cell communication between murine Zeb1 KO astrocytes and T cells (Fig. 4H), we decided to further investigate and validate how CXCL14 may modulate this interaction.

**Figure 6:**
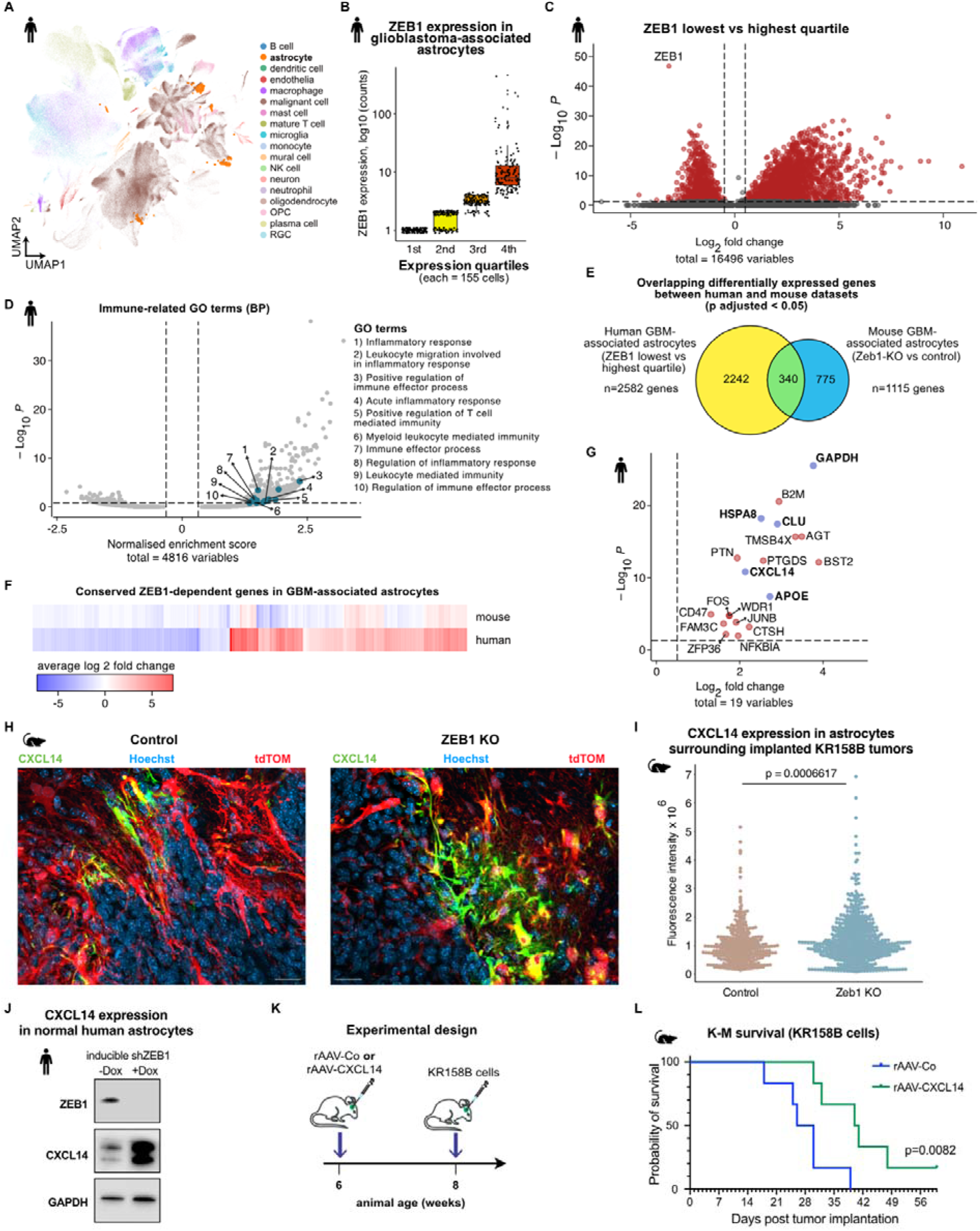
scRNAseq analysis of human GBM patient datasets reveals signatures of immune activation in GAA stratified for ZEB1 expression. **(A)** UMAP plot from combined human patient scRNAseq data (CELLxGENE, [29]), highlighting clusters of astrocytes. **(B)** Histogram showing ZEB1 expression in astrocyte cluster and indicating stratification of astrocytes into ZEB1-hi (top 25%ile) and ZEB1-lo (bottom 25%ile). **(C)** Volcano plot showing DEGs in astrocytes stratified for ZEB1 expression (P < 0.05, log_2_FC > 0.5). **(D)** GSEA enrichment plot for gene ontologies (GO BP) in astrocytes stratified for ZEB1 expression. Immune-related GO terms are highlighted in teal and key terms are labelled (P < 0.05, normalized enrichment score > 0.5). **(E)** Venn diagram showing overlapping DEGs between human astrocytes stratified for ZEB1 and mouse Zeb1 KO vs control astrocytes, corrected for false discovery. **(F)** Heat map showing overlapping ZEB1-dependent DEGs between human and mouse datasets. **(G)** Volcano plot showing overlapping ZEB1-dependent DEGs with unidirectional gene expression changes in both datasets (P < 0.05, log_2_FC > 0.5). Candidates involved in immune signatures identified in panel D are highlighted in blue. **(H)** Immunofluorescence imaging of CXCL14 in tissue sections from control (left) and Zeb1 KO (right) mice transplanted with KR158B GBM cells. Similar results were observed in 4 transplants per genotype. **(I)** Quantification of mean fluorescence intensity in mouse tissue sections (n=338 cells from 4 mice (control) and 569 cells from 4 mice (Zeb1 KO). **(J)** CXCL14 Western blot in normal human astrocytes transduced with control or ZEB1 knockdown vectors. Representative for n=3 repeat experiments. **(K)** Experimental design of CXCL14 overexpression in vivo. Adeno-associated viral vectors for CXCL14 or GFP expression were injected 2 weeks prior to intracranial implantation of KR158B GBM cells. **(L)** Kaplan-Meier survival analysis of CXCL14 overexpressing and control (GFP) mice transplanted with KR158B GBM cells (n=6 mice per genotype).

### Astrocytic CXCL14 is inversely regulated by ZEB1 and increases survival of tumor-bearing animals

Immunofluorescence staining for CXCL14 in brain sections from tumor-bearing control and Zeb1 KO mice (Fig. 6H) validated increased CXCL14 protein levels following *Zeb1* deletion (Fig. 6I). We further validated that ZEB1 knockdown in cultured normal human astrocytes increased CXCL14 protein expression (Fig. 6J). Finally, we injected recipient naïve C57BL6/J mice with adeno-associated viral vectors designed to deliver CXCL14 or empty virus as a control (GFP), followed by orthotopic transplantation of murine KR158B GBM cells (Fig. 6K). Targeted CXCL14 expression significantly increased survival of tumor-bearing mice compared to controls (Fig. 6L). Together our data support that ZEB1 inversely regulates CXCL14 in astrocytes that subsequently prevents efficient T-cell activation which in turn is pro-tumorigenic. However, ZEB1 loss or CXCL14 gain results in significant increase in survival of GBM-bearing mice via an improved T cell response and represents a potential novel treatment option to improve T-cell responses within the TME.

## Discussion

Astrocytes have emerged as important players in the GBM TME, and recent studies have highlighted how astrocyte-immune interactions promote an immunosuppressive environment and that targeting these interactions is a viable therapeutic strategy to improve survival in experimental GBM models [1, 2, 14, 15]. While it is evident that astrocytes secrete immunoregulatory cytokines that silence the anti-tumor immune response, how these processes are regulated in astrocytes remain unclear. We show that immunosuppressive actions are linked to astrocytic cell plasticity, and that blocking astrocytes from entering more stem-like cell states by deleting *Zeb1* disrupts immunosuppressive signaling. A previous study has shown that TRAIL+ astrocytes suppress T cell activation [2]. We did not find altered TRAIL expression following astrocytic *Zeb1* deletion indicating that increased immune activation following *Zeb1* loss may be independent of TRAIL. This supports GBM reprogramming GAA into a state of increased cell plasticity that promotes anti-inflammatory astrocyte-immune signaling; astrocytic deletion of *Zeb1* prevents this reprogramming and causes astrocytic secretion of pro-inflammatory cytokines that promote T cell recruitment and activation which results in immune clearance of tumors.

Astrocytes enter reactive cell states upon neural injury that are characterized by increased plasticity and more stem-like phenotypes [30]. For example, reactive astrocytes surrounding CNS lesions show increased expression of cytoskeletal markers that are expressed in neural stem cells, such as Nestin and Vimentin [3]. Reactive astrocytes can adopt different heterogeneous cell states that range from neuroprotective (anti-inflammatory) to neurotoxic (pro-inflammatory), yet the molecular drivers of these cell states are incompletely understood [31]. Of the two astrocyte clusters in our dataset, the astrocyte_2 cluster showed a stronger reduction in *Zeb1* expression, cell plasticity and stemness-associated signatures, as well as immune signatures. Our data support that ZEB1 is a transcriptional regulator of anti-inflammatory cell states in GAA and *Zeb1* deficiency results in lymphocyte recruitment and activation.

ZEB1 is a master regulator of cell plasticity and well-known for its functions on cancer stemness and epithelial-mesenchymal transition [32]. Beyond cell plasticity, ZEB1 also regulates DNA damage repair and immunosuppressive signaling molecules in tumor cells [32]. Recent evidence has further shown that ZEB1 is necessary for polarization of cancer-associated fibroblasts in colorectal cancer, where *Zeb1* deficiency increases cytokine production [20]. Our data suggest that ZEB1 has similar functions in astrocytes in GBM and that GAA share immunosuppressive traits with cancer-associated fibroblasts in other solid-tissue cancers.

We identify CXCL14 as a candidate mediator of immune function in the GBM TME and potential therapeutic target. Upregulation of CXCL14 in *Zeb1*-deficient astrocytes promotes recruitment and activation of T-cells, including cytotoxic lymphocytes. Notably, CXCL14 expression in human astrocytes with low abundance of ZEB1 correlates with GAA in Zeb1 KO mice. The convergence of single-cell ligand-receptor inference, flow cytometric validation of brain-localized T cell activation, and independent therapeutic benefit from CXCL14 delivery collectively support an astrocyte-T cell mechanism in our study. There is some controversy in the literature whether CXCL14 has oncogenic or tumor suppressive functions [33, 34] indicating that CXCL14 may have context- and cell-type-dependent functions. It is noteworthy that previous studies investigating CXCL14 effects in vivo have relied on immunocompromised models, with our data suggesting that CXCL14’s anti-tumorigenic effects via immune activation may be stronger that the pro-tumorigenic effects on GBM cell proliferation. Our work clarifies the functions of CXCL14 in an immunocompetent system and supports anti-tumor activity of CXCL14 which can convert a typically “cold” TME into an environment characterized by T cell recruitment. This supports the prospect of using CXCL14 as an immunotherapeutic agent for GBM [35].

In conclusion, our work demonstrates that cell plasticity in GAA promotes GBM immune evasion through anti-inflammatory signaling, which can be counteracted by CXCL14. Therefore, CXCL14 is a potential therapeutic candidate for reprogramming the TME that can restrict GBM growth and progression.

## Materials and Methods

### Cell culture

Murine GBM KR158B cells were provided by Dr Karlyne Reilly (NCI) and NPE-IE cells were provided by Prof Steven M. Pollard (University of Edinburgh). KR158B cells were cultured in N2 medium (Thermo Fisher Scientific) supplemented with Mycozap Plus (Lonza), 20ng/ml each EGF (Peprotech) and thermostabilized FGF2-G3 (Qkine) [36]. NPE-IE cells were cultured in N2/B27 medium (Thermo Fisher Scientific) supplemented with 1% penicillin/streptomycin solution (Thermo Fisher Scientific), 10ng/ml each EGF (Peprotech) and thermostabilized FGF2-G3 (Qkine) and 3 µg/ml Laminin (Sigma) [22]. Normal human astrocytes (ScienCell Research Laboratories) were cultured in astrocyte medium supplemented with 2% fetal bovine serum, 1% penicillin/streptomycin solution and 1% astrocyte growth supplement (all ScienCell Research Laboratories). Lentiviral transduction of GBM cells and normal human astrocytes with fluorescent reporters or doxycycline-inducible ZEB1 knockdown constructs was performed as described previously [37].

### Animal experiments

Animal care and handling and all procedures were performed according to the Federation of European Laboratory Animal Science Associations (FELASA) and institutional guidelines and approved by the UK home office (PP0188547 and PP9348382). Orthotopic xenografts were performed in female SCID mice as described previously [37]. Syngeneic transplants of murine GBM cells were performed in C57Bl6/J mice (Charles River) or in transgenic GLAST::CreERT^2^,Zeb1^f/f^,R26-loxP-Stop-loxP-tdTomato (Zeb1 KO) or GLAST::CreERT^2^,R26-loxP-Stop-loxP-tdTomato mice (control) [19, 38, 39] of both sexes. Intracranial tumor transplants were performed as described previously [37]. Depending on the experiment and cell line, 5×10^4^ – 2×10^5^ cells were stereotactically implanted in 5 µl of DMEM/F12. Mice were maintained under isoflurane anesthesia during procedures. Mice were monitored regularly for development of neurological symptoms and body weight loss. For histological analysis, animals were transcardially perfused with 4% paraformaldehyde and the brains removed and postfixed in 4% paraformaldehyde. For flow cytometry analysis, blood was removed via cardiac puncture and animals were transcardially perfused with PBS with the removed brains and blood processed for FACS. For scRNAseq, animals were transcardially perfused with PBS and the brains removed and tumor-containing regions microdissected using a fluorescent stereomicroscope. Four brains from each genotype were pooled and processed for single cell analysis using an OctoMACS dissociator and adult brain dissociation kits according to manufacturer’s instructions (Miltenyi). For viral delivery of CXCL14, animals were injected intracranially with 10^11^ vector genomes of recombinant adeno-associated viral vectors (rAAV5) expressing murine CXCL14 and a fluorescent reporter, or GFP as control. Two weeks after virus injection, animals were intracranially injected with 5×10^4^ KR158B cells as described above.

### Human tissue samples

Tissue samples from human GBM patients were obtained from the University Hospital Wales. Fixed tissue samples were cryoprotected, embedded in OCT (Thermo Fisher Scientific), and sectioned on a cryomicrotome (Leica) at a thickness of 100 µm. Free-floating tissue sections were stained as described [19].

### Immunofluorescence staining

Post-fixed brains were cryoprotected, embedded and sectioned as described [37]. Immunofluorescence staining was performed using standard protocols. A list of antibodies is provided in Supplementary Table 1.

### Protein Isolation and Western Blotting

Protein isolation, quantification, and Western blotting were performed as described [37]. GAPDH was used as loading control.

### Flow cytometry

Blood and brain derived leukocytes from control or Zeb1 KO mice were stained as described previously [40, 41]. Briefly, cells were stained in order of Zombie Aqua (BioLegend), then with anti-CD16/CD32 Fc-block (BioLegend), followed by anti-mouse conjugated monoclonal antibodies listed in Supplementary Table 1. Data were acquired using an Attune NxT flow cytometer (Thermo Fisher Scientific) and analyzed using FlowJo version 10 (FlowJo LLC).

### Image acquisition and analysis

Fluorescent images were acquired on a Zeiss LSM710 inverted confocal microscope using ZEN software or Olympus Slideview VS200 slide scanner using VS200 ASW software. Images were imported into ImageJ for analysis. For quantification of tumor size and invasion, we used the ‘analyze particles’ function in ImageJ on thresholded images (GFP channel) to measure the number of individual satellite tumors (proportional to invasion) and the area covered by the tumor (proportional to tumor size).

### Single cell RNA sequencing

For preparation of single cell libraries, cells from 2 mice per genotype were pooled and 40,000 cells per genotype were loaded onto a Chromium Next GEM chip G (10XGenomics). Libraries were prepared using 10XGenomics Library Construction Kit according to manufacturer’s instructions. RNA QC was performed using a Qubit and Tapestation. RNA sequencing was performed at Wales Gene Park using an Illumina NovaSeq 6000 and S1 flow cell. Fastq files were downloaded from the sequencing provider and run through the nf-core/scrnaseq pipeline (v.2.4.0; [42]) where they were aligned to the mouse genome reference (mm10) using 10X Genomics’ CellRanger (v.6.1.1) *count* tool. Count matrices were then imported into R (v.4.3.2) and processed using the Seurat analysis pipeline (v5.0.2, [43]). Briefly, samples were read into R and filtered for a minimum of 3 cells, 200 minimum features, then following plotting of quality control (QC) metrics, were further filtered for > 200 and < 5000 nFeature_RNA (unique features), and < 10% mitochondrial gene expression per cell. Samples were then log normalized, cell cycle scores were calculated, and variable features were selected (n = 3000). Samples were then integrated using Seurat’s *FindIntegrationAnchors* and *IntegrateData* functions using the previously selected variable features. The integrated dataset was then scaled with variable regression (nCount_RNA, percent.MT, G2M score, and S score) and principal component analysis was performed with 30 principal components (PC’s). Neighbors were then found, using 30 PC’s, followed by clustering with a resolution of 0.1. UMAP analysis was then performed using default parameters and data visualized using the scCustomize package (v.2.0.1, [44]). Clusters were annotated using a combination of methods: cluster differentially expressed genes were calculated using the *FindAllMarkers* function and analyzed in Enrichr, and known cell type markers were visualized using the *AddModuleScore* function and via feature plots, violin plots, and heatmaps. Gene set enrichment analysis (GSEA) was then performed on each cluster using the *singleseqgset* package (v.0.1.2.9000, [45]) and visualized using the *pheatmap* package (v.1.0.12, [46]). GSEA and visualization of differentially expressed gene sets between genotypes was performed using the genekitr package (v1.2.8, [47]), using Benjamini-Hochberg method for statistical correction and cutoff values for p of 0.05 and q of 0.15. Ligand-Receptor analysis was performed using the LIANA package (v.0.1.12) with default parameters [28]. Copy number variation analysis was performed on the samples via the inferCNV package (v.1.3.3, [48]), using default parameters and the Gencode murine GRCm38 gene order file. Velocity analysis was performed using the *scVelo* python package (v.0.2.5, [49]) with default parameters. Loom files for scVelo analysis were generated using the *Velocyto* package (v.0.17, [50]) and the Ensemble GRCm38 murine gene annotation file. To check Zeb1 expression across publicly available datasets, we used the CELLxGENE API in Python following their tutorial (v.1.17.0, census v.2.1.0, [29]).

### Data analysis and statistical testing

Imaging and survival data analysis was performed in GraphPad Prism version 10. Normality distribution was evaluated using the Shapiro-Wilk test. Two-tailed Welsh’s t-test was used for statistical comparison of normally distributed data and two-tailed Mann-Whitney test was used where data was not normally distributed. Kaplan-Meier survival analysis was performed using Log-Rank test. Statistical analysis of scRNAseq data was performed in R (version 4.3.2) using appropriate packages. No statistical methods were used to predetermine sample sizes and sample sizes were chosen based on those reported in previous publications [2, 18]. Data collection experiments were not randomized, and investigators were not blinded. In all analyses, p values <0.05 were deemed statistically significant. Statistical significance was indicated as follows: \**P* < 0.05; \*\**P* < 0.01; \*\*\**P* < 0.001; \*\*\*\**P* < 0.0001.

## Acknowledgements

This work was supported by the Medical Research Council (MR/X018318/1, awarded to FAS), and The Brain Tumour Charity (GN-000754 awarded to MC).

## Author contributions

**MC**: Investigation, Writing – original draft, Writing – review and editing, Funding acquisition. **AG**: Formal analysis, Data curation, Writing – review and editing. **AB**: Investigation, Writing – review and editing. **DS**: Investigation. **SK**: Investigation. **NS**: Formal analysis. **BG**: Methodology, Investigation, Visualization. **VE**: Methodology, Investigation. **FAS**: Writing – original draft, Writing – review and editing, Visualization, Conceptualization, Supervision, Funding acquisition.

## Data availability

Datasets from this study will be available at NCBI’s Gene Expression Omnibus at the time of publication.

## Supplemental information

**Figure S1:**
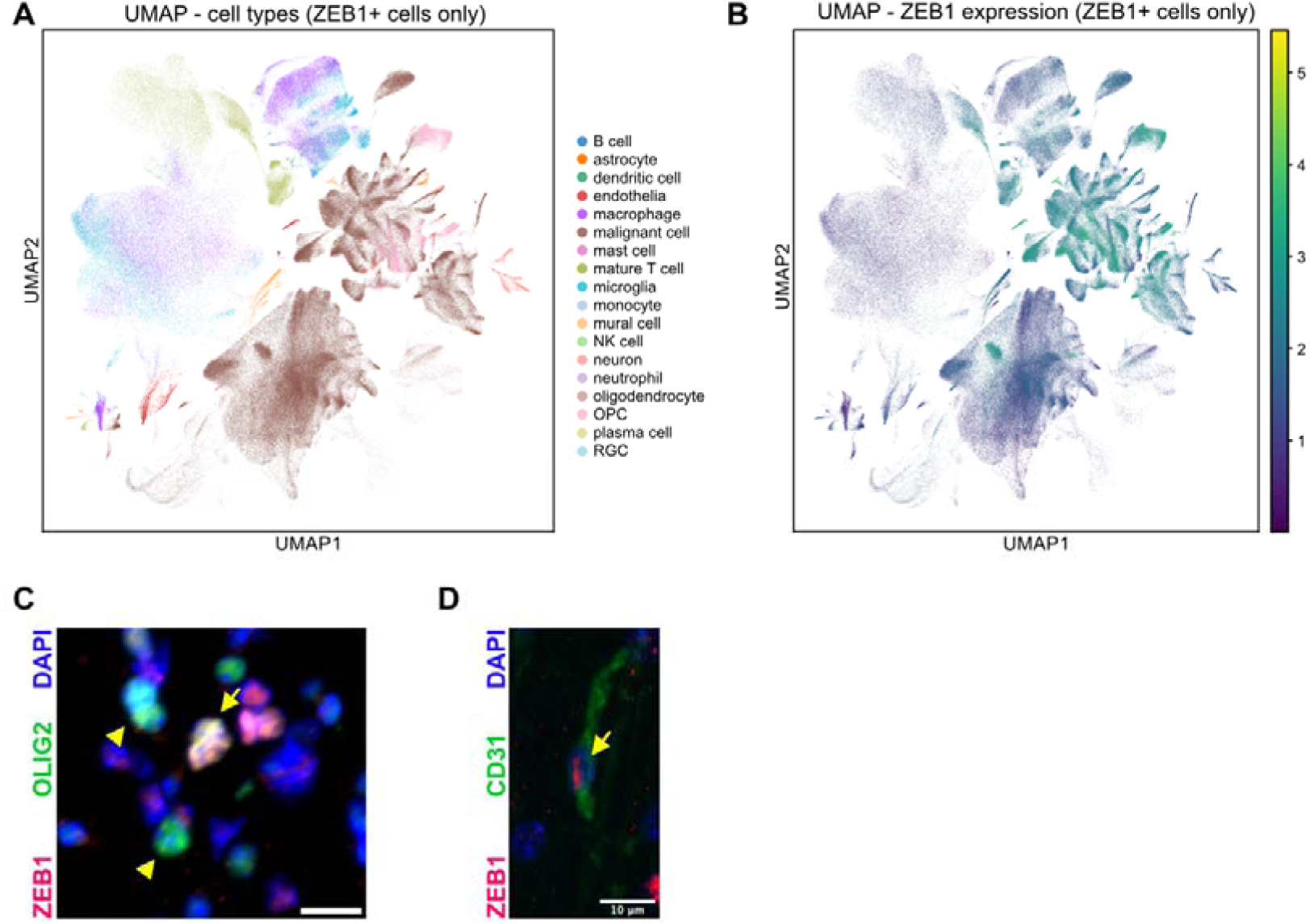
**(A)** UMAP plot of ZEB1-expressing cells in the Extended GBmap dataset, labelled by cell type. **(B)** UMAP plot of ZEB1 expression in the Extended GBmap dataset (cells with zero read counts for ZEB1 removed). **(C)** Some cells in the adult mouse cortex co-express ZEB1 and the oligodendrocyte precursor cell marker OLIG2 (arrow), whereas some oligodendrocyte precursor cells are negative for ZEB1 (arrowheads). **(D)** Expression of ZEB1 in CD31-positive vascular endothelial cells. Scale bars 10 µm. Images in C,D representative for three independent mice.

**Figure S2:**
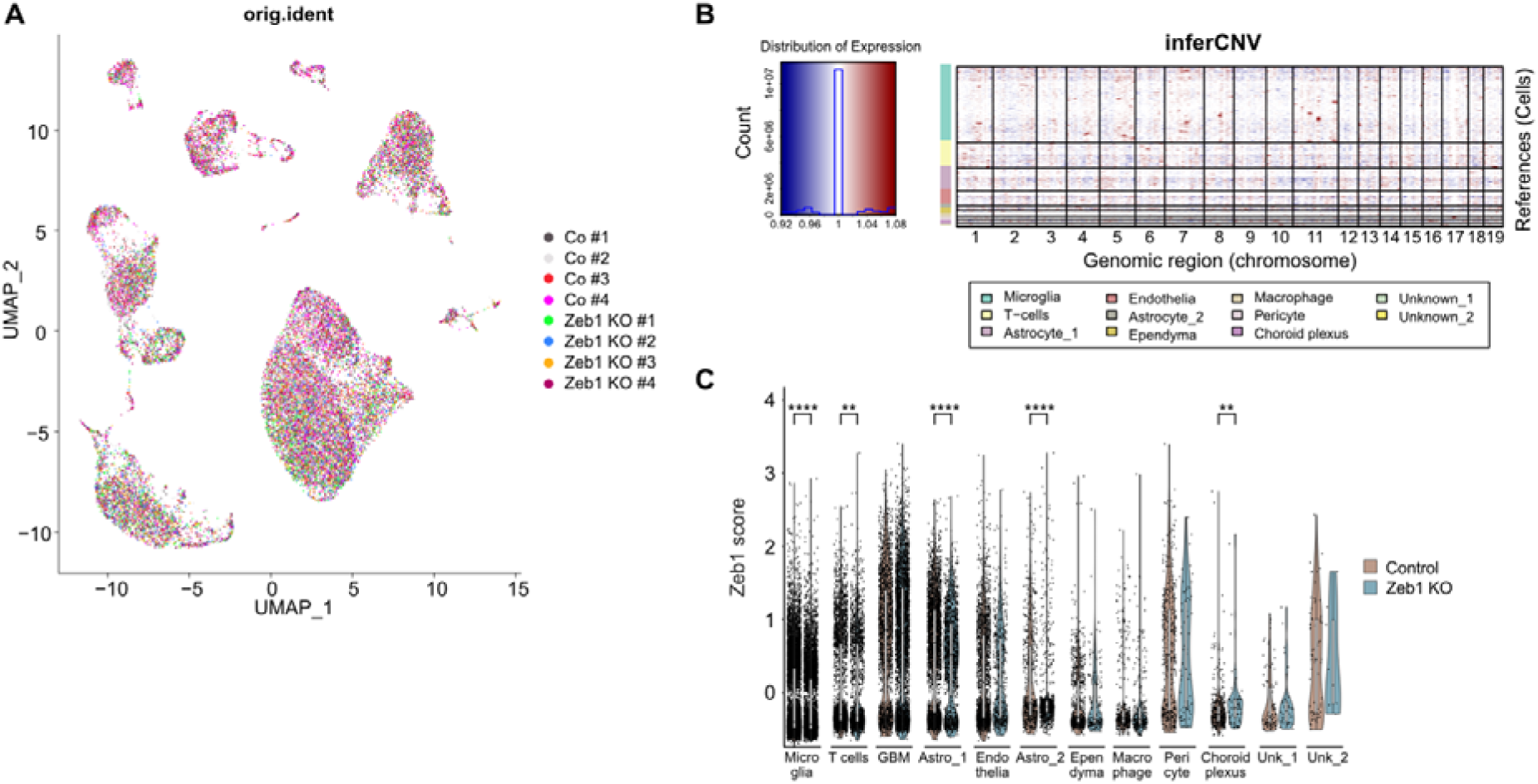
**(A)** UMAP plot of Seurat clusters labelled by sample showing no significant contribution to individual clusters by single samples. **(B)** InferCNV plot showing no significant genomic alterations in non-malignant cell clusters. **(C)** Violin plots showing *Zeb1* expression across Seurat clusters. Note that *Zeb1* expression is lowest in the astrocyte_2 cluster.

**Figure S3:**
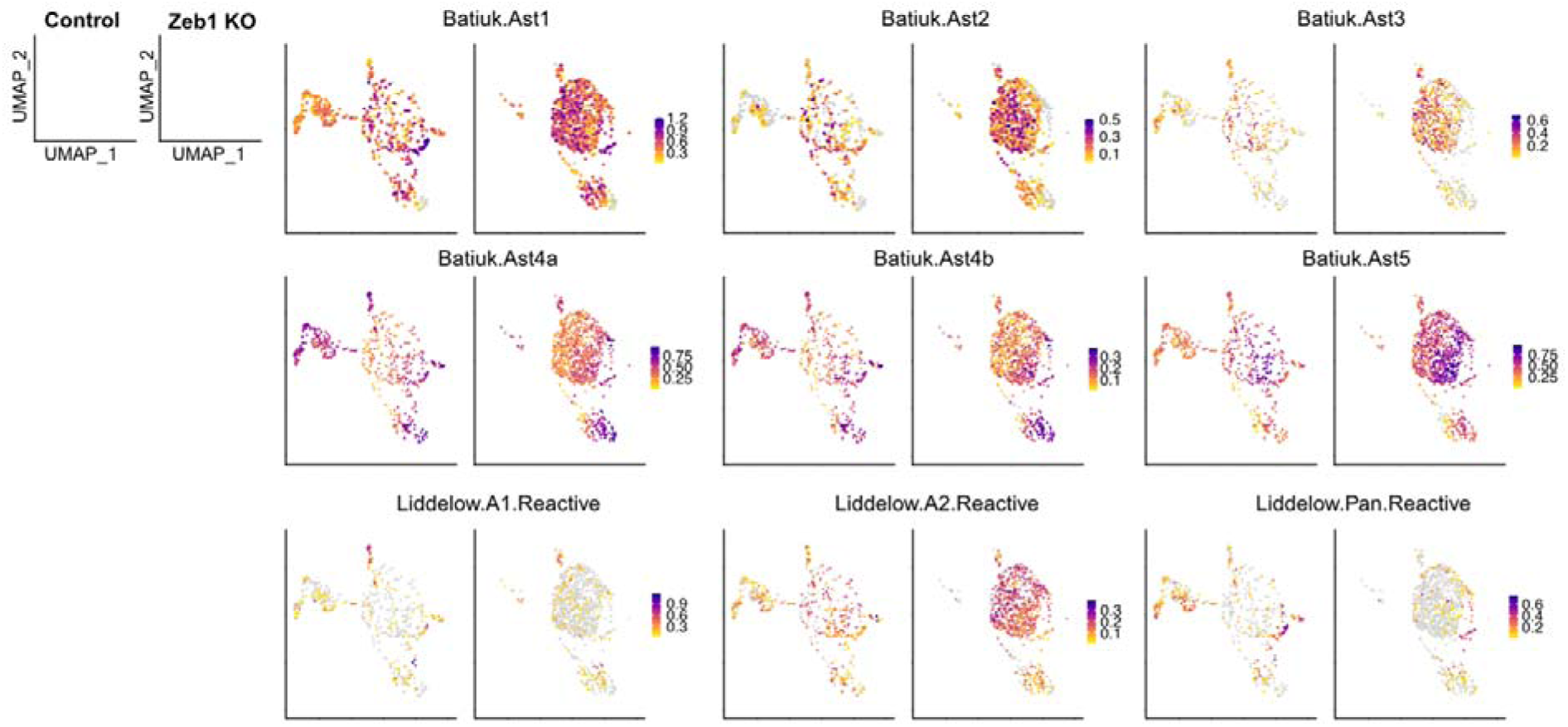
UMAP plots of the astrocyte_2 cluster only, comparing expression levels of region-specific astrocyte subtypes [23] and reactive astrocyte subtypes [24] between control and Zeb1 KO mice.

**Figure S4:**
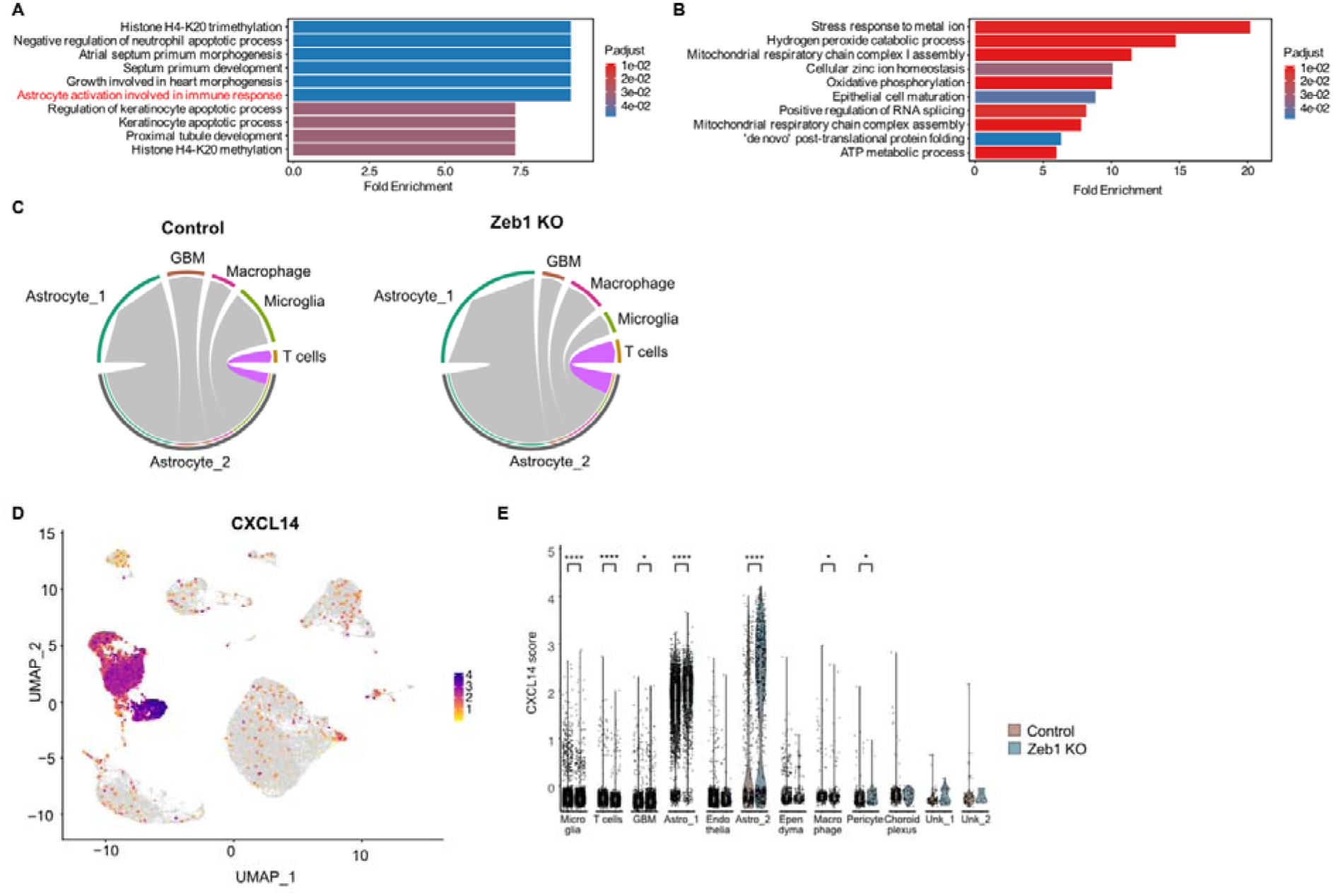
**(A)** Top 10 gene ontologies (BP) over-represented in the astrocyte_2 cluster. Immune-related ontologies highlighted in red. **(B)** Top 10 gene ontologies (BP) over-represented in the astrocyte_1 cluster. **(C)** Chord diagrams of cell-cell interactions between astrocyte_2 (sender) and other astrocyte, GBM, or immune cell clusters (recipients) in the GBM TME of Control or Zeb1 KO mice. Signaling to T cells is highlighted in purple. **(D)** UMAP plot showing expression of CXCL14 is almost exclusively confined to the astrocyte_1 and astrocyte_2 clusters. **(E)** Violin plots showing expression of CXCL14 across all clusters in control and Zeb1 KO samples.

**Figure S5:**
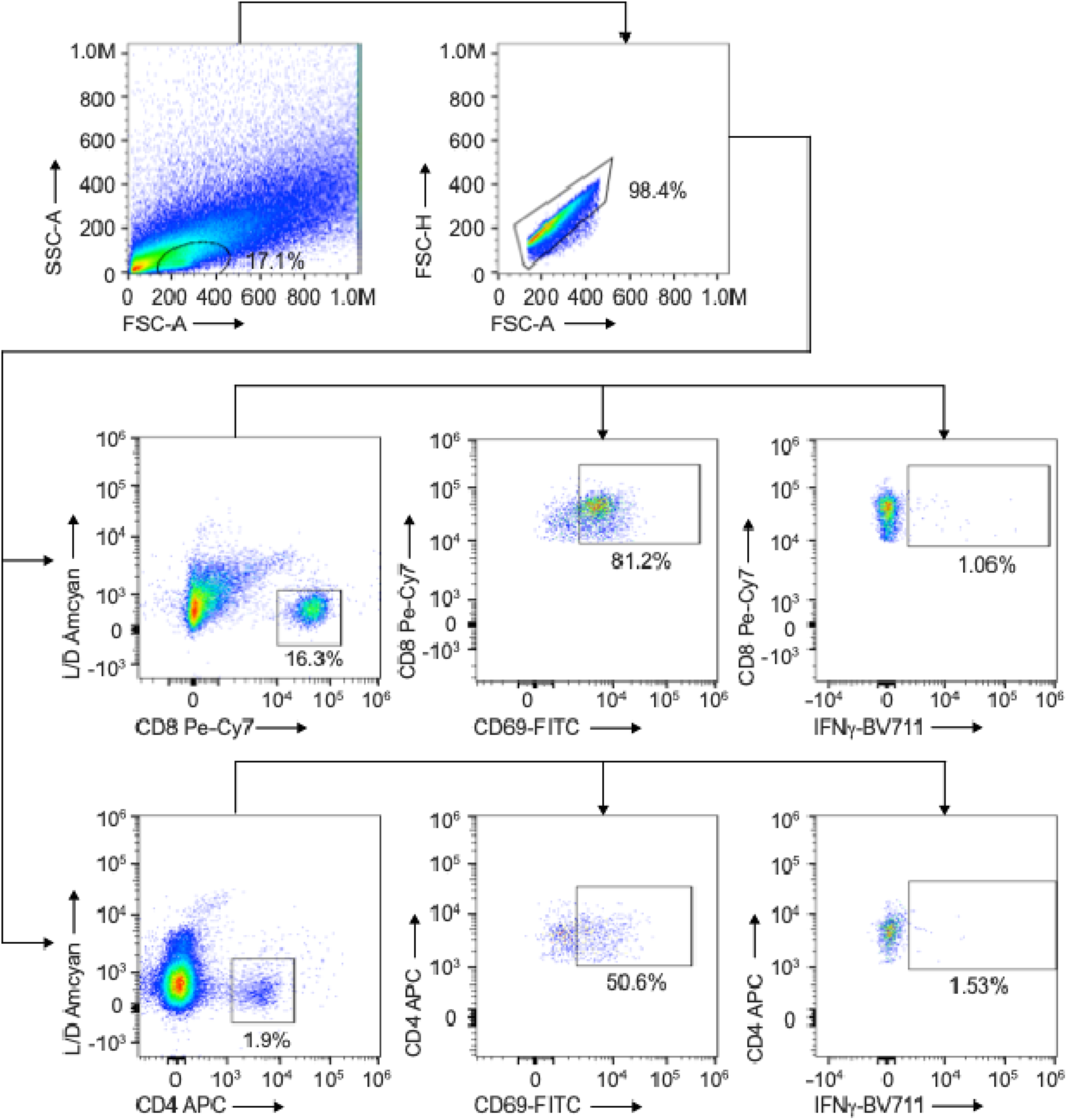
Flow cytometry gating strategy. Leukocytes were harvested from brains and were strained as described in the materials and methods. Samples were run and thresholds were set for FSC-A vs SSC-A to assign gates for debris exclusion. This population was sub-gated using FSC-H vs FSC-A for singlet cell identification. This population was sub-gated using expression of Live/Dead fixable stain with either CD4 or CD8 where Live/Dead negative and either CD4 or CD8 positive cells were identified as live cells. The live CD4 or CD8 cells were sub-gated to identify the expression of either CD69 or IFN-gamma where gates and percentages indicate positive cells. FMO controls were used for each expression marker (excluding Live/Dead) to define set gate boundaries. Cells were acquired using an Attune NxT flow cytometer and analysed using FlowJo.

**Table S1:** List of antibodies used in this study.

| Primary Antibody | Species | Dilution | Supplier | Catalogue number |
| --- | --- | --- | --- | --- |
| bIII-tubulin (TuJ) | Mouse | 1:500 | Promega | G7121 |
| CD4-APC (clone RM4-5) | Rat | 1:100 | BioLegend | 100516 |
| CD8-PE-Cy7 (clone 53.6.7) | Rat | 1:100 | BioLegend | 100722 |
| CD31 | Goat | 1:100 | R&D Systems | AF3628 |
| CD44-PerCP (clone IM7) | Rat | 1:50 | BioLegend | 103036 |
| CD45-BV605 (clone 30-F11) | Rat | 1:100 | BioLegend | 103139 |
| CD62L-PE (clone MEL-14) | Rat | 1:100 | BioLegend | 104408 |
| CD69-FITC (clone H1.2F3) | Armenian Hamster | 1:100 | BioLegend | 104506 |
| CNPase | Mouse | 1:250 | Merck | MAB326 |
| CXCL14 | Rabbit | 1:100 | Novus | NBP1-31398 |
| GFAP | Chicken | 1:1000 | Encor | CPCA-GFAP |
| GFAP | Mouse | 1:250 | Sigma | G6171 |
| GFP | Sheep | 1:2000 | Novus | NBP100-62622-0 |
| Hoechst 33342 | - | 1:500 | Thermo Fisher | H3570 |
| IFNg-BV711 (clone XMG1.2) | Rat | 1:100 | BioLegend | 505835 |
| Nestin (human) | Mouse | 1:500 | Encor | MCA-4D11-100 |
| OLIG2 | Rabbit | 1:500 | Merck | AB9610 |
| TNFa-AF700 (clone MP6-XT22) | Mouse | 1:100 | BioLegend | 506338 |
| ZEB1 | Rabbit | 1:500 | Sigma | HPA027524 |

